# Gene expression associated with white syndromes in a reef building coral, *Acropora hyacinthus*

**DOI:** 10.1101/012211

**Authors:** R. M. Wright, G. V. Aglyamova, E. Meyer, M. V. Matz

## Abstract

****Background**:** Corals are capable of launching diverse immune defenses at the site of direct contact with pathogens, but the molecular mechanisms of this activity and the colony-wide effects of such stressors remain poorly understood. Here we compared gene expression profiles in eight healthy *Acropora hyacinthus* colonies against eight colonies exhibiting tissue loss commonly associated with white syndromes, all collected from a natural reef environment near Palau. Two types of tissues were sampled from diseased corals: visibly affected and apparently healthy.

****Results**:** Tag-based RNA-Seq followed by weighted gene co-expression network analysis identified groups of co-regulated differentially expressed genes between all health states (disease lesion, apparently healthy tissues of diseased colonies, and fully healthy). Differences between healthy and diseased tissues indicate activation of several innate immunity and tissue repair pathways accompanied by reduced calcification and the switch towards metabolic reliance on stored lipids. Unaffected parts of diseased colonies, although displaying a trend towards these changes, were not significantly different from fully healthy samples. Still, network analysis identified a group of genes, suggestive of altered immunity state, that were specifically up-regulated in unaffected parts of diseased colonies.

****Conclusions**:** Similarity of fully healthy samples to apparently healthy parts of diseased colonies indicates that systemic effects of white syndromes on *A. hyacinthus* are weak, which implies that the coral colony is largely able to sustain its physiological performance despite disease. The genes specifically up-regulated in unaffected parts of diseased colonies, instead of being the consequence of disease, might be related to the originally higher susceptibility of these colonies to naturally occurring white syndromes.

## Background

Increasing rates of disease have contributed greatly to global coral population declines over the last few decades [1, 2]. The broadly defined “white syndromes” in Indo-Pacific regions, characterized in the field by tissue loss resulting in exposure of the coral skeleton, have been attributed to *Vibrio* spp. [3], a genus of bacteria involved in several coral diseases [4–8]. Other reports find no evidence of pathogenic bacteria in diseased corals [9] and instead blame stress-triggered programmed cell death for the manifestation of symptoms [10]. These conflicting conclusions, drawn mostly from culturing assays and histological observations, are further confounded by insufficient knowledge of the cnidarian immune response.

Corals, like all invertebrates, rely entirely on innate immunity for protection from invading pathogens. Features of innate immunity in corals include physical barriers [11], molecular pattern recognition [12], secretion of antimicrobial macromolecules [13], and cellular responses (*e.g.*, phagocytosis) [14–16]. Recent efforts to characterize those features of immunity using various cnidarian genome and transcriptome sequence databases have identified putative components of coral stress management and immune response pathways by homology with better-studied organisms. The *Acropora digitifera* genome project revealed striking differences in innate immunity complexity in corals compared to a closely related cnidarian, *Nematostella vectensis* [17]. Whereas the *N. vectensis* genome encodes only a single Toll/Toll-like receptor (TLR), the *A. digitifera* genome included at least four TLRs, along with other related immune signaling molecules. Miller *et al.* reported the presence of TLR signaling components, including adaptor proteins that link that cascade with other signaling events, in the expressed sequence tag library of another acroporid coral, *A. millepora* [12]. Together these elements suggest an ability of corals to respond to pathogen-associated molecular patterns (PAMPs) via TLR recognition and integrate that signal to cellular responses such as inflammation and apoptosis. Toll/TLR signaling can activate NF-κB transcription factor that, upon nuclear localization, up-regulates transcription of immune response genes. In corals, the identities of those response genes and the roles they play remain unclear. Some suggested immune response genes include lectins, complement c3 and apextrin, proteins involved in non-self recognition, aggregation and cell lysis respectively [18]. Metabolism and calcification genes demonstrated differential expression in addition to immunity genes in *A. millepora* challenged with bacterial and viral immunogens, providing a more comprehensive picture of cellular events during an acute infection [19]. Global RNA-sequencing of *A. cervicornis* displaying signs of White Band Disease (WBD) revealed that disease significantly affected the expression of genes involved in immune processes and apoptosis [20]. The up-regulation of phagocytic cell surface receptors and reactive oxygen species (ROS) producing enzymes suggested that the phagocytosis and degradation of damaged cells drives the WBD response in corals. These coral sequencing projects and experimental immune challenges have provided conclusive evidence that corals are capable of launching defensive responses upon direct contact with pathogens. A coral’s ability to communicate the recognition of that pathogen along the colony, however, is less understood. Coral polyps utilize a gastrovascular system lined with flagellated gastrodermal cells to transport organic products and zooxanthellae within the colony [21]. These channels are used to allocate energetic resources to areas that need them most, such as fast-growing branch tips [22–24] and wounded regions [25]. Radiolabeled carbon accumulation experiments have shown that corals preferentially direct energetic resources *towards* physically damaged regions [25] but *away* from disease-induced lesions [26]. These findings suggest that healthy coral tissues might possess means to detect and respond to an advancing disease lesion, but it is still unclear what the physiological consequences of this action might be. Here we examine the gene expression profiles of *A. hyacinthus* displaying white syndromes (Figure 1) to determine the molecular consequences of the diseased condition. White syndromes advance along a colony in a way such that a distinct lesion forms between affected and unaffected tissues. Tissues ahead of the lesion are presumably healthy, while tissues at the lesion boundary are actively sloughing cells in response to infection. We compared gene expression profiles among three health states: affected tissues (diseased, “D”), apparently healthy tissues from diseased colonies (“**a**head of the **l**esion”, “AL”), and tissues from completely unaffected colonies (healthy, “H”). Comparing healthy regions of diseased colonies to completely disease-free individuals provided an opportunity to look for expression patterns that might indicate a colony-wide systemic effect of infection and/or disease susceptibility. We used tag-based RNA-Seq [27] followed by weighted gene correlation network analysis [28] to achieve systems-level insight into molecular responses to chronic disease in corals.

**Figure 1:**
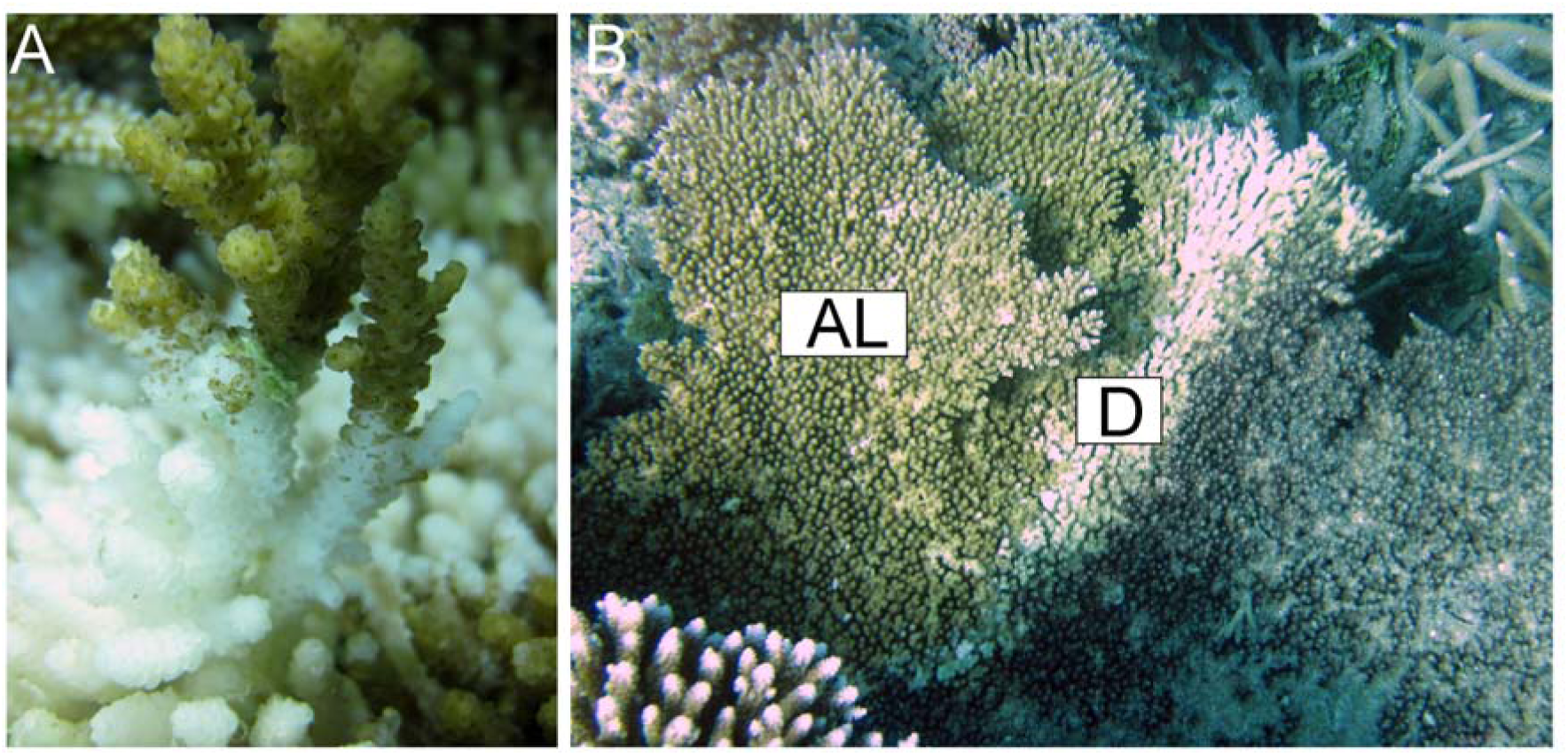
White syndrome in *A.hyacinthus* sampled in this study. (A) Close-up of the lesion area. (B) Position of sampled locations in diseased colonies: “D” – diseased, “AL” – ahead of the lesion. Tissues were also sampled from completely healthy individuals (“H”, not pictured). Photo credit: Carly Kenkel.

## Results

### Differential Gene Expression between Health States

Sequencing yielded an average of 6,367,219 reads per sample. An average of 19.5% of these remained after filtering and of these, an average 31.45% mapped to the transcriptome. A total of 44,701 isogroups (clusters of contigs representing the same gene, from here on referred to as “genes”) were detected. Reads were converted to unique transcript counts by removing PCR duplicates, yielding an average of 156,650 counts per sample (Additional File 1). A generalized linear model with contrasts between all three tissues detected differentially expressed genes between health states (Additional File 2). The disease-healthy contrast yielded 646 DEGs passing a Benjamini-Hochberg FDR cutoff of 10%. The disease-AL contrast yielded 333 DEGs passing an FDR cutoff of 10%. No genes passed the 10% FDR cutoff for the healthy-AL contrast. Between all contrasts, a total of 757 genes passed the FDR cutoff of 10%.

Principal coordinate analysis of the variance-stabilized data for all genes revealed expression differences mainly between disease (D) and the other two health states (Figure 2A). PCoA using the 3827 isogroups with an unadjusted p-value of less than 0.05 for any contrast revealed more differences between health states, but a significant overlap in expression of healthy and AL tissues remains (Figure 2B).

**Figure 2:**
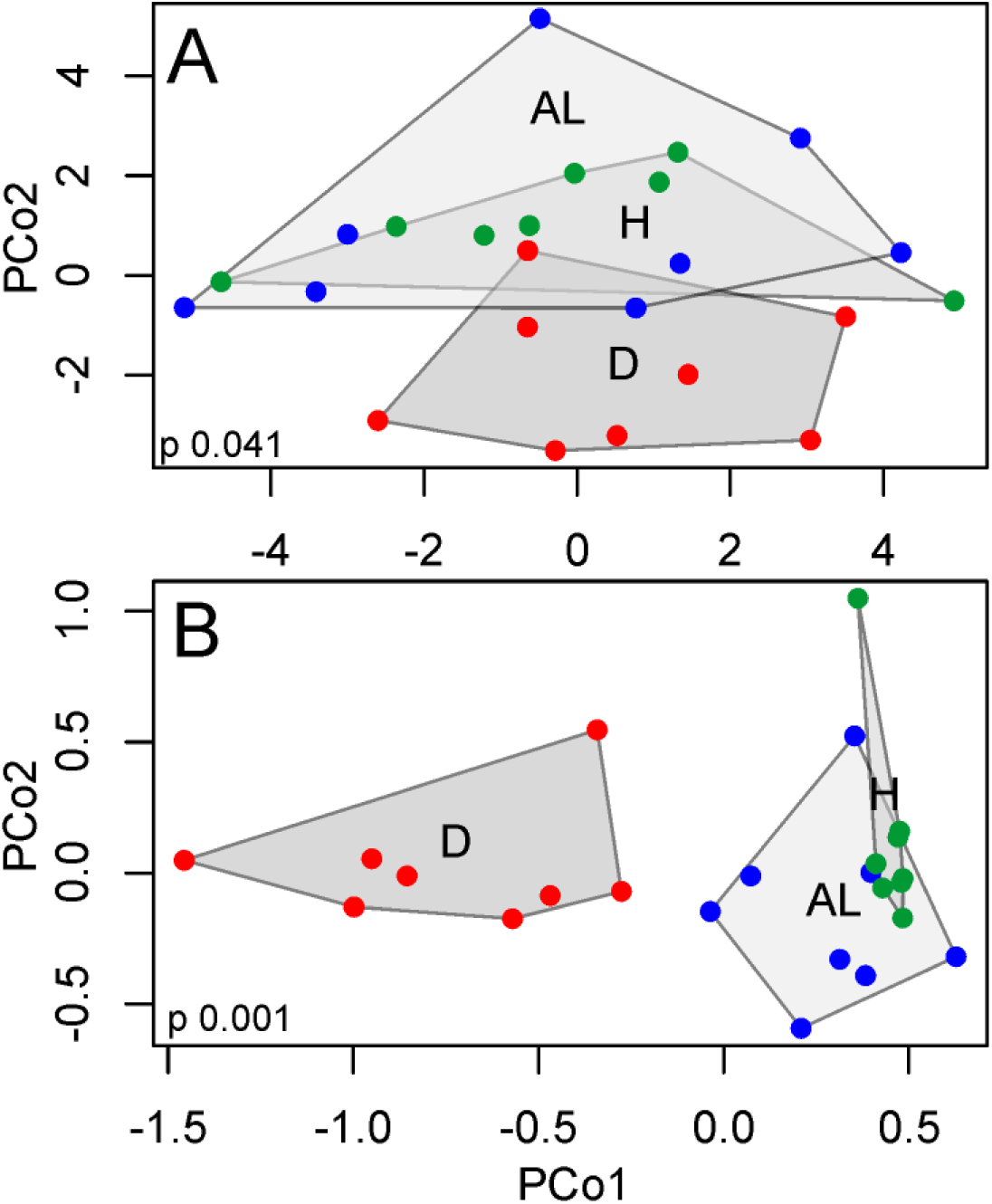
Principal coordinate analysis clusters samples by health state. Samples cluster by the presence of disease symptoms (D vs. AL and H) when all genes are included in the PCoA (A). Differences between health states become more evident when PCoA is performed on DEGs (unadjusted p-value < 0.05) only (B).

### Gene Ontology (GO) Enrichment

Functional enrichments between all three contrasts allow a general examination of the molecular functions and biological processes being differentially regulated between health states. The enriched groups of both the disease-healthy and disease-AL contrasts were largely identical (Figure 3 and Additional File 3). Ribosomal proteins, oxidative stress responses, and translation factor activity were up-regulated in diseased tissues compared to both AL and healthy tissues. Likewise, receptor activity, regulation of biological quality, and extracellular matrix components (collagens) were down-regulated in diseased tissues compared to both healthier states. No GO terms were significantly enriched (FDR 10%) for the healthy-AL contrast.

**Figure 3:**
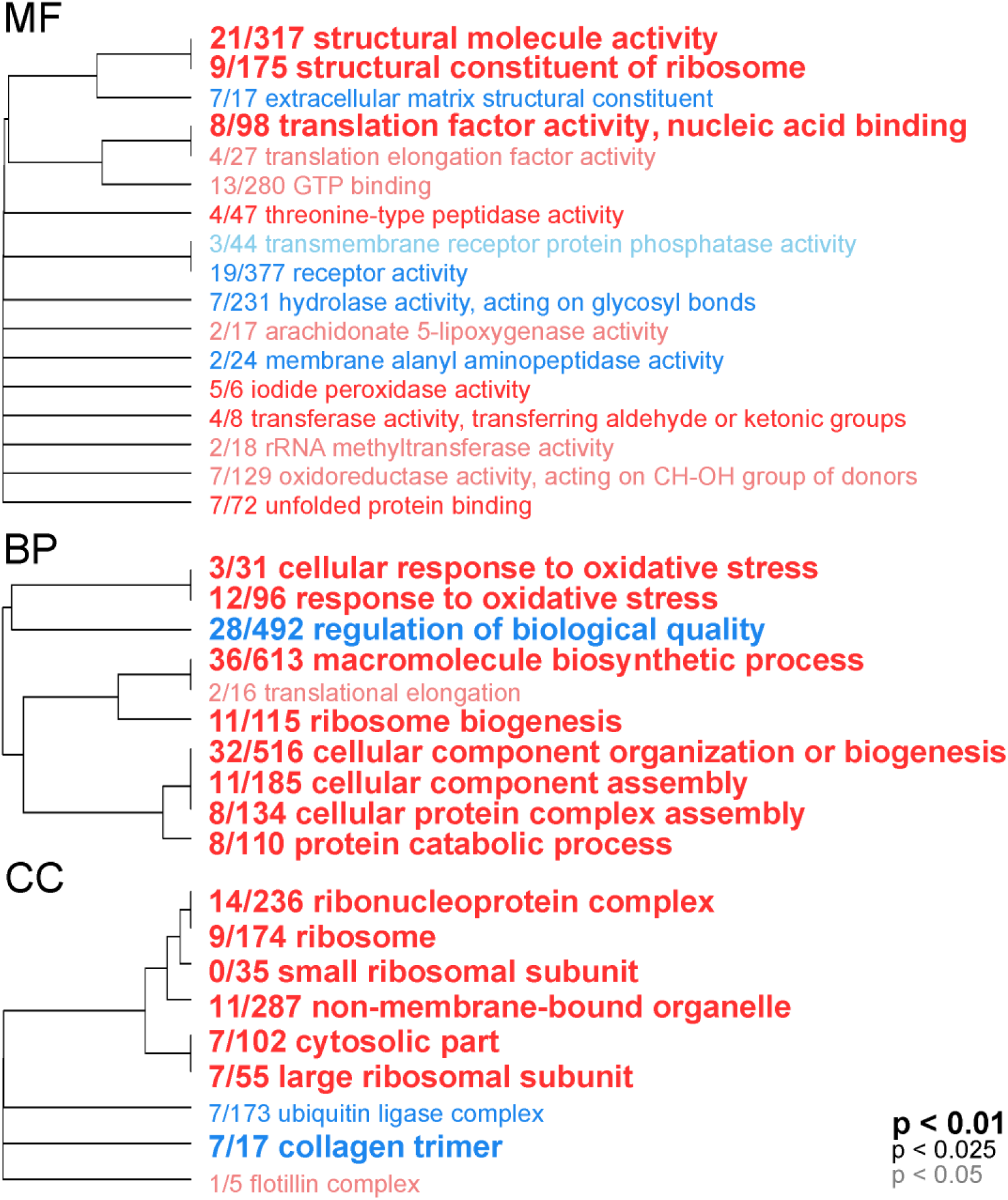
Gene ontology categories enriched by genes up-regulated (red) or down-regulated (blue) in diseased compared to fully healthy samples, summarized by molecular function (MF), biological process (BP), and cellular component (CC). The size of the font indicates the significance of the term as indicated by the inset key. The fraction preceding the GO term indicates the number of genes annotated with the term that pass an unadjusted p-value threshold of 0.05. The trees indicate sharing of genes among GO categories (the categories with no branch length between them are subsets of each other).

### Gene Expression Analysis by Contrast

Gene expression heatmaps were constructed to show the relative expression patterns of the top most significant DEGs for each contrast (Figure 4A and Additional Files 4,5). Complete lists of all annotated DEGs with log fold changes for each contrast can be found in Additional Files 6-8.

**Figure 4:**
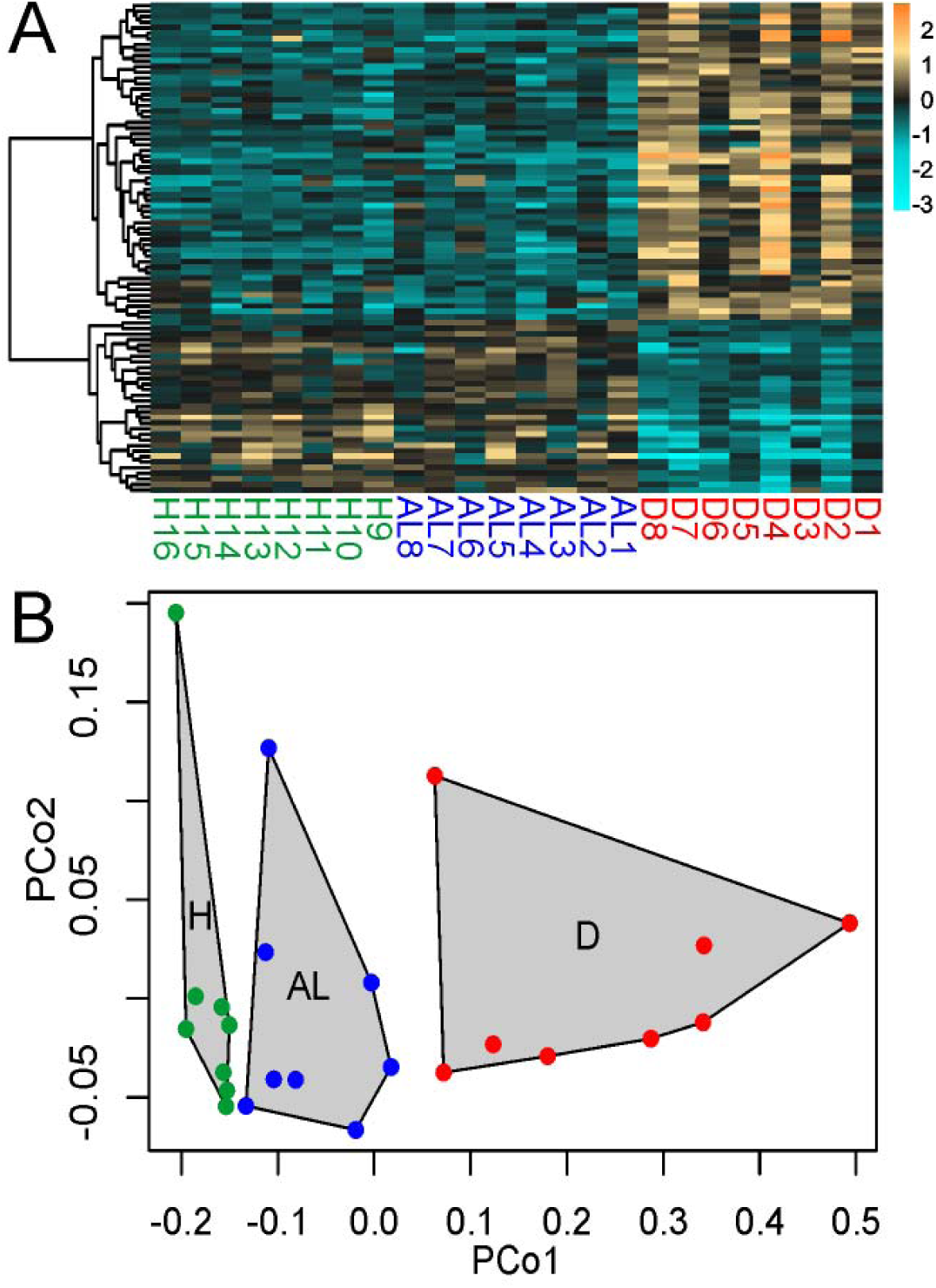
Expression of DEGs significant for disease-healthy contrast among health states. (A) Heatmap for top DEGs (FDR = 0.01). Rows are genes, columns are samples ordered as in the bottom panel: ahead-of-lesion (AL), healthy (H), and diseased (D). The color scale is in log_2_ (fold change relative to the gene’s mean). The tree is a hierarchical clustering of genes based on Pearson’s correlation of their expression across samples. (B) Principal coordinate analysis of all DEGs at 10% FDR for disease-healthy contrast.

#### Diseased vs. Healthy

Genes found to be up-regulated (Benjamini-Hochberg FDR < 0.01) in diseased tissues compared to healthy corals include key members of the oxidative stress response in corals (*e.g.*, catalases and peroxidases) and pentose phosphate metabolism (transketolase, transaldolase, and 6-phosphogluconate dehydrogenase). Both proteinases (astacin and cathepsin L) and protease inhibitors (alpha-macroglobulin and serine proteinase inhibitor Ku-type) were up-regulated in diseased tissues. Two of the genes annotated as C-type lectin, a carbohydrate-binding protein, and malate synthase, a key enzyme of the glyoxylate cycle, were also up-regulated in symptomatic tissues. Down-regulated genes (Benjamini-Hochberg FDR < 0.01) include those encoding extracellular matrix constituents (collagens, heparin sulfate proteoglycans) and carbonic anhydrase, a key enzyme in coral skeletal deposition. Red fluorescent protein was also down-regulated in diseased tissues, a hallmark of the coral stress response [29–33].

#### AL vs. Diseased and Healthy

The expression differences between diseased and AL tissues within a colony paralleled the expression differences between diseased tissues and healthy corals. At the same significance threshold (Benjamini-Hochberg FDR < 0.01), almost the exact same top-candidate genes were identified (Additional Files 4,5). No differentially expressed genes passed the Benjamini-Hochberg 10% FDR when comparing the AL tissues and healthy corals.

To better quantify the behavior of disease-responsive genes in AL samples, the expression of genes passing a 10% FDR cutoff for the disease-healthy comparison was studied using principal coordinate analysis. Tukey’s test based on the first principal coordinate values revealed highly significant differences between H and D as well as between AL and D (P < 0.001 in both cases). Although AL samples appeared to be intermediate between H and D samples (Figure 4B), their scores along the first principal coordinate axis were not significantly different from healthy samples (P = 0.11).

### Correlation Between Gene Network Modules and Health States

A total of 6737 DEGs with unadjusted p-value < 0.1 were input into WGCNA for network analysis. A sample network was constructed to identify outlying samples with a standardized connectivity score of less than -2.5. One sample (diseased individual “4”) was identified as an outlier and removed from subsequent analysis (Additional File 9). Twelve unique modules, assigned arbitrarily color labels, remained after merging highly correlated modules. Of these twelve modules, eight were highly correlated to a single coral individual and one (grey) is reserved to contain genes that do not fall into any coexpression module. The remaining three modules were highly correlated with the health states (Additional File 10). Since we assembled these modules using a signed network, the sign of the correlation is equivalent to the direction of expression change with respect to the trait. For example, a module that is significantly negatively correlated to diseased corals contains genes that are down-regulated in that state.

The eigengene of the dark green module (1155 genes) was strongly correlated with diseased-healthy contrast (Pearson’s *R*^2^ = 0.83, P_cor_ = 1e-6, Additional File 10). The genes within this module are up-regulated in diseased tissues and down-regulated in healthy tissues, while tending to be down-regulated in AL samples. Conversely, the turquoise module (669 genes) was up-regulated in healthy tissues and down-regulated in diseased tissues (Pearson’s *R*^2^ = -0.83, P_cor_ = 9e-7, Additional File 10). Expression of these two modules in AL samples demonstrated similar direction of change as in healthy tissues, although the change was not statistically significant. Notably, one module was identified (green, 661 genes) that was significantly up-regulated in AL (Pearson’s *R*^2^ = 0.64, P_cor_ = 0.001) and (to a lesser extent) down-regulated in diseased tissues (Pearson’s *R*^2^ = -0.44, P_cor_ = 0.04), while remaining unchanged in healthy tissues (Additional File 10).

### Within Module Gene Expression Analysis

Hierarchically clustered gene expression heatmaps were constructed to show the relative expression patterns of the genes within each module that best represent the module and show significant correlations to the health state, based on module membership and gene significance values (Figure 5 and Additional File 11).

**Figure 5:**
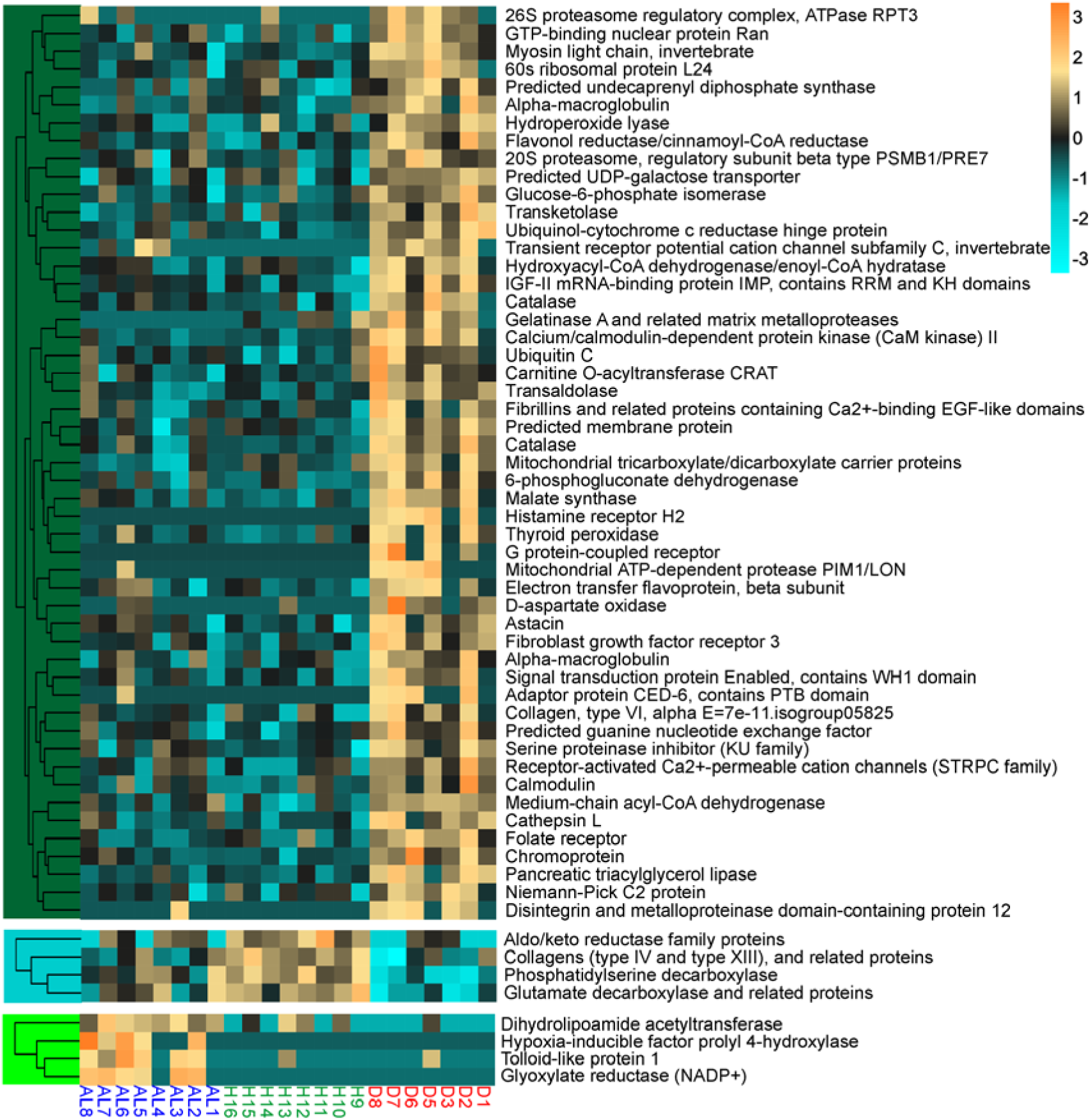
Gene expression heatmaps of annotated DEGs with a module membership and gene significance score greater than 0.6. Rows are genes, columns are samples ordered as in the bottom panel: ahead-of-lesion (AL), healthy (H), and diseased (D). The color scale is in log_2_ (fold change relative to the gene’s mean). The trees are hierarchical clustering of genes based on Pearson’s correlation of their expression across samples. The color block of the trees indicates the module to which these genes belong (dark green, turquoise, and green).

## Discussion

The major pattern of variation in gene expression was between asymptomatic (healthy and AL) and diseased (D) corals (Figure 2). There were no statistically significant differences in gene expression between healthy colonies (H) and asymptomatic parts of diseased colonies (AL), suggesting that these states are physiologically similar. The genes that are differentially regulated between diseased and healthy corals show a subtle trend towards disease-like gene expression in asymptomatic tissues of diseased colonies (Figure 4). This trend is, however, not statistically significant, indicating that white syndromes have little effect on the physiology of the unaffected portion of *A. hyacinthus* colony. Gene co-expression network analysis revealed groups of genes co-regulated with respect to each of the three states, including a group of genes specifically up-regulated in AL samples.

### Diseased tissues up regulate immune response elements

Innate immunity provides immediate protection against non-self and responds to physical injury. Three general steps are involved in an innate immune response: detection, defense activation, and effector responses to neutralize the threat. Tissues sampled from the lesion of disease progression in corals exhibiting white syndromes have enhanced expression of genes involved in each of these three immune response phases. C-type lectins act as pattern recognition receptors to activate pathogen elimination through phagocytosis in invertebrates [34]. Cnidarian genomes encode *c-type lectin* genes with highly variable substrate regions, leading to hypotheses that these proteins recognize a large variety of pathogens [35]. In *A. millepora*, mannose-binding C-type lectins have been shown to respond immediately following an immune challenge (only 45 minutes after lipopolysaccharide injection in [36]), but show no significant response at later time points [37]. The up-regulation of C-type lectins in tissues at the lesion may suggest that these tissues have very recently become infected. The second phase of an immune response prepares targets for elimination via antimicrobial peptide synthesis and immune cell activation. While this experiment did not discover any differentially regulated antimicrobial peptides, we do detect the activation of immune activating proteins C4, alpha-macroglobulin and CD109 in diseased tissues. The lectin pathway of immune activation is triggered by lectins binding a pathogen-associated molecule and results in the activation complement component factor C4 and C3 [38]. These complement factors, along with alpha-macroglobulin and CD109, tag pathogens and secreted proteases for elimination. In the final phase of an innate immune response, foreign organisms are engulfed and destroyed by phagocytic immune cells. Lysosomes within these phagocytic cells contain proteins capable of degrading engulfed material via the production of reactive oxygen species (ROS) or proteolytic enzymes, such as cathepsins. The up-regulation of cathepsin L in diseased tissue may be a consequence of such phagocytic activity. The up-regulation of immune-related transcripts in diseased corals is consistent with previous studies of both naturally occurring disease and experimental pathogen challenges [19, 20, 39]. Just like the rest of genes exhibiting H-D difference, these responses are confined to the symptomatic regions of the coral. One possible explanation of this fact is that the immunity-related gene expression changes are elicited by direct contact with a pathogen rather than a systemic signal throughout the colony.

### Switch to lipid-based metabolism in diseased tissues

Transcripts involving lipid and carbohydrate metabolism (triacylglycerol lipase, phosphoenolpyruvate carboxykinase), including glyoxylate cycle metabolism (malate synthase), were up-regulated in diseased tissues compared to asymptomatic tissues. The differential regulation of these metabolic genes suggests that diseased corals may be utilizing stored energy reserves more than healthy corals. Fatty acids derived from stored lipids are oxidized by the beta-oxidation pathway and release acetyl-CoA to enter the citric acid cycle. Both carbons in one molecule of acetyl-CoA are consumed during the decarboxylation steps of the citric acid cycle and energy is released. The glyoxylate cycle is an alternative route through the citric acid cycle that allows organisms to thrive on two-carbon sources by catalyzing the conversion of acetyl-CoA to malate and succinate via a glyoxylate intermediate, bypassing the decarboxylation steps in the citric acid cycle [40]. These four carbon compounds contribute to the energetic requirements of the cell and serve as building blocks for cellular components, fulfilling all the same necessities of the citric acid cycle without the need to replenish oxaloacetate from the diet. One of the most significantly up-regulated transcripts in diseased coral tissues was malate synthase, one of the key enzymes of the glyoxylate cycle along with isocitrate lyase. Glyoxylate cycle enzymes are rare throughout the animal kingdom, but bioinformatic analyses suggest they exist in cnidarians [41]. As additional support for a potential role of glyoxylate cycle metabolism in coral stress responses, glyoxylate cycle transcripts were up-regulated in *A. palmata* larvae subjected to thermal stress [42]. In higher plants, glyoxylate enzymes are active when the cell is switching from photosynthetic production of sugars to scavenging pathways from stored and structural lipids, as in starvation and/or senescence [43]. In corals, this metabolic shift might indicate a decline in shared energy reserves with zooxanthellae, presumably due to stress-induced symbiont loss.

### Oxidative stress response genes are up-regulated in diseased tissues

Reactive oxygen species (ROS) are produced as a consequence of fatty acid oxidation. The up-regulation of antioxidants that protect the cell from these harmful byproducts corals could be a consequence of increased fatty acid metabolism. This explanation coincides well with the observed up-regulation of lipid metabolism and antioxidant (catalase, peroxidase) transcripts in diseased corals. The production of ROS is also a fundamental element of the innate immune response. While ROS are capable of neutralizing phagocytized pathogens, the harm they cause to the host must be countered if an organism is to withstand its own immune response. Catalases and peroxidases capable of hydrolyzing harmful peroxides provide a mechanism of such self-protection. The upregulation of oxidative stress response genes is well characterized in corals experiencing thermal stress [44], physical stress [45], and infectious disease [20].

### Matrix metalloproteinases are up-regulated in diseased tissues

Stony corals are subject to many potential sources of physical injury such as predators [46], boring organisms [47] and storms [48, 49]. The tissue regeneration mechanisms employed by corals that have sustained a physical injury are common to wound-healing processes across metazoans [50]. One of these steps involves restructuring of the extracellular matrix to encourage tissue regeneration. Matrix metalloproteinases (MMPs) are a group of enzymes capable of such activities and have been shown to play a direct role in wound repair in Hydra [51]. In addition, MMPs act on pro-inflammatory cytokines to direct inflammation due to wounding and innate immune responses to pathogens (reviewed in [52]). The up-regulation of MMPs in response parasitic protists in a gorgonian coral suggests that these proteins are active in the immune response of cnidarians [53]. Astacin and gelatinase have matrix metalloproteinase activities and were up-regulated specifically in affected coral tissues. Additionally, a protease inhibitor alpha-macroglobulin was up-regulated, which is a vital component of the innate immune response that inactivates bacterial secreted proteases, thus compromising their virulence [54].

### Calcification genes are down-regulated in diseased tissues

Calcification rates in reef-building corals are sensitive to several environmental variables such as light, pH, and temperature [1, 55–57]. While this experiment did not directly measure coral calcification, the identification of DEGs with functions in biomineralization suggests that disease negatively impacts coral skeletal deposition. Both general extracellular matrix structural components and coral-specific calcification functions were differentially regulated in diseased tissues compared to asymptomatic tissues. Coral biomineralization is directed by an extracellular skeletal organic matrix comprised of secreted glycosylated proteins [58]. These proteins include collagens and negatively charged macromolecules (like chondroitin sulfate proteoglycans) that bind calcium ions to aid in crystal formation [59]. Several collagens and a protein with high similarity to nematogalectin, a collagen family protein that forms a major structural component in Hydra nematocyst tubules [60], were down-regulated in diseased tissues. The down-regulation of these genes in diseased tissues suggests a weakening of the coral skeletal organic matrix and thus a diminished capacity for biomineral deposition. The potential impact of disease on coral skeletal growth is most clearly revealed by the downregulation of carbonic anhydrase, an enzyme that plays a fundamental role in mediating bicarbonate supplies for calcification in scleractinian corals [61–63].

### AL-specific gene expression: a systemic response to disease or factors contributing to disease susceptibility?

Genes that are specifically regulated in apparently healthy tissues of diseased colonies could represent systemic (*i.e*., colony-wide) response to disease. However, there is another possible interpretation: since expression of these genes is correlated with natural appearance of disease, it might signify disease susceptibility rather than disease effects. A recent study employing similar a sampling scheme to investigate transcriptomic effects of yellow band disease (YBD) in *Orcibella faveolata* [39] found that expression in asymptomatic regions of diseased colonies was intermediate between completely healthy corals and diseased tissue, which fits well with the systemic response interpretation. In our study the AL expression was most similar to the healthy state (Figure 2 and 4A), although genes differentially regulated between diseased and healthy states demonstrated a non-significant trend towards intermediate expression in AL samples (Figure 4B). The difference between the two studies could potentially be explained by unequal levels of colony integration between *Orbicella* and *Acropora* ([64] and references therein), which could affect the extent of the systemic signaling and/or spread of the pathogen throughout the colony. The co-expression network analysis revealed a sizeable (661 genes) module that was up-regulated in the AL state compared to D and H states (Additional File 10). Among the genes most strongly associated with this module were the genes coding for the immunity-related Tolloid-like protein and the hypoxia inducible factor prolyl 4-hydroylase (HIF-P4H, Figure 5). Up-regulation of HIF-P4H suggests that healthy tissues of diseased colonies might be experiencing hypoxic conditions [65] [66]. Notably, HIF-P4H has also been shown to modulate immune responses by modifying the kinase responsible for releasing NF-κB from its inhibitor [67]. Up-regulation of these genes in healthy parts of diseased colonies might therefore be a sign of altered immunity state potentially explaining higher disease susceptibility of the affected colonies in nature.

## Conclusions

Our gene expression analysis identified several immune, repair, and metabolic molecular pathways expressed in coral regions affected with white syndromes. In contrast to *Orbicella faveolata*, *A. hyacinthus* does not show pronounced propagation of these responses to regions of the colony not visibly affected by disease, suggesting that the effect of chronic white syndromes on colony-wide *A. hyacinthus* physiology is small. Instead, asymptomatic regions of diseased colonies show gene expression signatures potentially related to higher disease susceptibility of the affected coral individuals. Further studies of natural disease-associated gene expression will contribute towards the development of diagnostic tools to predict and manage coral disease outbreaks.

## Abbreviations

AL: Ahead-of-the-lesion

DEG: Differentially expressed genes

FDR: False discovery rate

FOG: Fuzzy orthologous groups

GO: Gene ontology

HIF-P4H: Hypoxia-inducible factor prolyl 4-hydroxylase

KEGG: Kyoto encyclopedia of genes and genomes

KOG: EuKaryotic orthologous groups

MMP: Matrix metalloproteinase

NF-κB: Nuclear factor kappa-light-chain-enhancer of activated B cells

PAMP: Pathogen-associated molecular pattern

ROS: Reactive oxygen species

TLR: Toll-like receptor

WBD: White band disease

WGCNA: Weighted gene correlation network analysis

YBD: Yellow band disease

## Methods

### Sampling

Coral fragments from 16 colonies of *A*. *hyacinthus* were sampled in the spring of 2011 along the eastern coast of Palau (7° 18.738’ N, 134° 30.423’ E) and immediately stored in RNAlater (Ambion). Eight of these colonies were visibly affected with white syndromes (Figure 1). Colonies exhibiting diffuse tissue loss along a lesion of apparently healthy tissue directly adjacent to exposed white skeleton in accordance with [68] were sampled. Colonies displayed no obvious signs of predation. The remaining eight colonies were completely symptom-free (designated healthy, H). From the eight affected colonies, coral fragments were sampled from both the lesion interface between diseased and healthy tissues (diseased, D) and areas well ahead of the lesion (AL, Figure 1B). AL coral fragments were sampled from approximately midway between the lesion boundary and the edge of the colony. AL tissues are unlikely to be in direct contact with pathogen since previous studies have demonstrated declines in pathogens only ∼1 cm in advance of a white syndromes lesion [69].

### Transcriptome Assembly and Annotation

The *A. hyacinthus* transcriptome has been generated from 5-day aposymbiotic larvae as described previously [70]. It was annotated based on two resources: the proteome of the starlet sea anemone *Nematostella vectensis* [71] and in-depth annotations of the *Acropora digitifera* proteome [72]. Based on manual verifications of a subset of *A. digitifera* annotations, they were pre-filtered to include only protein sequences longer than 60 amino acids with the annotation assigned based on the listed e-value = 1e-20 or better. The GO, KEGG (“Kyoto Encyclopedia of Genes and Genomes”), KOG (“euKaryotic orthologous groups”), and gene name annotations were transferred to an *A. hyacinthus* contig if the contig matched one or both of these two resources with e-value = 1e-4 or better in blastx [73]. The GO and KOG annotations assigned to genes that were denoted FOG (“fuzzy orthologous group”, [74]) in the *N. vectensis* data were removed, since such genes encode proteins with common domains and cannot be functionally annotated based on homology alone. The annotated *A. hyacinthus* transcriptome has been released for unrestricted use prior to this publication,

http://www.bio.utexas.edu/research/matz_lab/matzlab/Data.html.

### Tag-based RNA-Seq

Libraries were prepared following [27] and sequenced using Applied Biosystems SOLiD v.3 platform. Read trimming, quality filtering, mapping, and conversion to per-gene counts was performed as described previously [27] with one modification: reads mapping to the same starting coordinate in the reference and aligning with 100% identity along the full length of the shorter read were discarded as potential PCR duplicates. The current step-by-step library preparation protocol as well as bioinformatic pipeline are available at https://sourceforge.net/projects/tag-based-rnaseq/; note however that the current version of the tag-based RNA-seq method utilizes advanced procedure for PCR duplicates removal based on degenerate tags incorporated during cDNA synthesis, and assumes sequencing on the Illumina HiSeq instrument.

### Identification of Differentially Expressed Genes (DEGs)

All statistical analyses were performed using R3.1.1 [75]. DEGs were identified using a generalized linear model implemented by the R package DESeq2 [76]. No outlying samples were detected by the arrayQualityMetrics package [77]. DESeq2 performed automatic independent filtering to remove lowly abundant transcripts and maximize the rate of DEG discovery post multiple testing correction at an alpha of 0.1. P-values for significance of contrasts between all three health states were generated based on Wald statistics and were adjusted for multiple testing using the false discovery rate method [78]. The contrasts resulted in tables including adjusted and unadjusted p-values and log_2_ fold changes that were used in downstream analyses.

### Gene Coexpression Network Analysis

A weighted gene correlation network analysis (WGCNA, [28]) was used to identify groups of co-regulated genes in an unsupervised way. Genes with an unadjusted p-value < 0.1 for any of the three contrasts as determined by the generalized linear model testing for the effect of health state were input into WGCNA. A sample network was constructed to identify outlying samples with a standardized connectivity score of less than -2.5 [79]. A signed gene co-expression network was constructed with a soft threshold power of 24. Groups of co-regulated genes (modules) correlated with each other with the Pearson correlation coefficient 0.42 or better were merged. The eigengenes of the resulting modules (the first principal component of the expression matrix corresponding to the genes included in the module) were correlated with health states (H, AL, or D).

### Assessing the Robustness of the Analysis

Low quality of the SOLiD sequencing resulted in low number of reads mapped, raising concerns about the reliability of the data. In tag-based RNA-seq, unlike standard RNA-seq, every count represents an observation of an independent transcript. Thus low counts could still provide sufficient quantitative information about transcript abundances. In addition, high level of biological replication (n=8 per group) in our experiment should have compensated for the low counts within each replicate to a certain degree. To confirm that low counts did not result in inflated false discovery rate, we have simulated a series of count datasets based on the empirical per-gene total counts and coefficients of variation across samples as well as empirical sample size factors, which included no effect of experimental conditions. Analysis of these simulated datasets recovered nearly identical sample size factors and highly similar dispersion estimates as in real data (Additional File 12). When these simulated datasets were analyzed with DESeq2 using the same models as real data, at most four genes passed the 10% Benjamini-Hochberg false discovery rate (FDR) cutoff for each contrast. Compared to 646 genes passing the FDR 10% cutoff for the disease-healthy comparison and 333 genes passing the same cutoff for the disease-AL comparison in the real dataset this is much less than 10%, indicating that the real data analysis was conservative. The simulation-based p-value cutoff achieving the empirical 10% FDR (Additional File 13A-C) would have yielded 1.05-1.5X more DEGs for these comparisons than the Benjamini-Hochberg procedure. Notably, the Benjamini-Hochberg correction did not yield any DEGs for the healthy-AL comparison, and accordingly, the DEG discovery rate in this comparison was even slightly lower than the simulation-based false discovery rate (Additional File 13C), indicating that for this comparison DESeq2 analysis did not provide sufficient power. To keep the analysis conservative, we chose to report DESeq2-based DEGs discovered using the Benjamini-Hochberg procedure. The procedure for inferring empirical FDR based on simulations described above has been implemented in the R package empiricalFDR.DESeq2, hosted within the Comprehensive R Archive Network (CRAN).

WGCNA analysis provides one additional confirmation that the observed gene expression differences are driven by biological factors rather than stochasticity. WGCNA constructs gene co-expression modules from their correlation pattern across samples without using the information about how the samples are distributed among experimental conditions. The fact that *post hoc* the module eigengenes correlate strongly with coral condition (Additional File 10) indicates that the gene expression patterns in the data truly reflect the biological processes related to disease. To verify this, we shuffled the condition designations among samples and indeed observed that the correlations with co-expression modules disappeared (Additional File 14).

### Principal Coordinate Analysis

Principal coordinate analysis to visualize clustering of gene expression between health states was performed using the “adegenet” package [80] using variance stabilized data for all genes and subsequently with only candidate differentially expressed genes (unadjusted p-value < 0.05), based on Manhattan distances which correspond to the sum of absolute log-fold changes across all genes. Effects of the three health states (“D”, “H”, and “AL”) were calculated using the multivariate analysis of variance function “adonis” of the R package “vegan” [81]. Tukey’s tests between specific health states based on the values of the first principal coordinate were performed using function TukeyHSD in R.

### Functional Summaries

Gene ontology enrichment analysis was performed using adaptive clustering of GO categories and Mann-Whitney U tests [82] based on ranking of signed log p-values (GO-MWU, https://sourceforge.net/projects/go-mwu/). Gene expression heatmaps with hierarchical clustering of expression profiles were created with the pheatmap package in R [83].

## Authors Contributions

GVA prepared the libraries for sequencing, EM assembled the transcriptome, RMW conducted the tag-based RNA-seq data analysis. MVM supervised the study, annotated the transcriptome, and performed simulations. MVM and RMW wrote the manuscript.

## Acknowledgements

We thank C Kenkel, D Stump, and C Palmer for sample collections and photographs. Funding for this study was provided by National Science Foundation grant DEB-1054766 to MVM.

